# Attention-dependent coupling with forebrain and brainstem neuromodulatory nuclei changes across the lifespan

**DOI:** 10.1101/2023.09.29.560190

**Authors:** Nicholas G. Cicero, Elizabeth Riley, Khena M. Swallow, Eve De Rosa, Adam Anderson

## Abstract

Attentional states continuously reflect the predictability and uncertainty in one’s environment having important consequences for learning and memory. Beyond well known cortical contributions, rapid shifts in attention are hypothesized to also originate from deep nuclei, such as the basal forebrain (BF) and locus coeruleus (LC) neuromodulatory systems. These systems are also the first to change with aging. Here we characterized the interplay between these systems and their regulation of afferent targets – the hippocampus (HPC) and posterior cingulate cortex (PCC) – across the lifespan. To examine the role of attentional salience on task-dependent functional connectivity, we used a target-distractor go/no go task presented during functional MRI. In younger adults, BF coupling with the HPC, and LC coupling with the PCC, increased with behavioral relevance (targets vs distractors). Although the strength and presence of significant regional coupling changed in middle age, the most striking change in network connectivity was in old age, such that in older adults BF and LC coupling with their cortical afferents was largely absent and replaced by stronger interconnectivity between LC-BF nuclei. Overall rapid changes in attention related to behavioral relevance revealed distinct roles of subcortical neuromodulatory systems. The pronounced changes in functional network architecture across the lifespan suggest a decrease in these distinct roles, with deafferentation of cholinergic and noradrenergic systems associated with a shift towards mutual support during attention guided to external stimuli.

**Significance statement:** Changes in attentional control across the lifespan may originate from cortical control networks or subcortical neuromodulatory systems, which are the first sites of age-related neuropathology. In young adults, we demonstrated functional coupling of the basal forebrain with the hippocampus, and locus coeruleus with the posterior cingulate cortex varies with task relevance. This coupling changed in middle age and most strikingly in older adults. In old age, task-dependent coupling between the locus coeruleus and basal forebrain was the predominant connection remaining within the observed network. Older adults exhibit reduced subcortical-cortical connectivity, consistent with a relative neuromodulatory deafferentation, replaced by subcortical-subcortical deep nuclear connectivity. This alteration in noradrenergic and cholinergic signaling has important implications for attention and memory formation and neurocognitive aging.

## Introduction

To produce adaptive behavior, the brain balances the prioritization of goal-relevant information with the need to respond to changing environmental demands (Shine, 2019). Depending on the situation, systems that process task-relevant information may predominate or give way to systems that promote shifts in cognitive states (Sarter et al., 2001; Corbetta et al., 2008). For example, when driving towards a traffic light, a yellow light could signal a switch from maintaining a steady speed to assessing if you need to slow down. The flexibility to dynamically shift attentional states is vital for cognitive functioning (Chun & Turk-Browne, 2007).

Subcortical neuromodulatory systems are the foundation of short-term shifts in the tuning of flexible brain networks towards objects of attention (Berridge & Waterhouse, 2003; Aston-Jones & Cohen, 2005). With distributed processes throughout the brain, the basal forebrain (BF) and locus coeruleus (LC) release neuromodulators, adjusting the responsivity of neurons without causing action potentials. The LC releases norepinephrine (NE) (Berridge & Waterhouse, 2003; Aston-Jones & Cohen, 2005) and the BF, which consists of four regions (Ch1-3 = medial septum (MS); Ch4 = Nucleus Basilis of Meynert (nbM)), releases acetylcholine (ACh). Both have prominent innervations in the posterior cingulate cortex (PCC) and hippocampus (HPC), which are involved in spatiotemporal representations of memory and internally-driven cognition, respectively (Levitt & Moore, 1978; Mesulam et al., 1983; Detari, Sembra, & Rasmusson, 1997; Espana & Berridge, 2006; Markello et al., 2018). HPC and PCC have significant functional connectivity with these neuromodulatory regions (Jacobs et al., 2018; Markello et al., 2018; Turker et al., 2021). The release of NE and ACh thus contributes to the PCC and HPC’s respective roles in cognition.

The LC and BF nuclei not only project to the HPC and PCC but are themselves strongly interconnected. The LC is the primary source of noradrenergic innervation to the nbM (Espana & Berridge, 2006) and show significant functional connectivity (Turker et al., 2021). NE and ACh have complementary roles in computing uncertainty in the environment, such that ACh signals expected uncertainty, suppressing expectation-driven information in environments with predictability (Yu & Dayan, 2005). In contrast, NE signals unexpected uncertainty in which sensory information violates top-down expectations. Additionally, NE flattens low-dimensional energy landscapes of cortical dynamics, reducing the difficulty of switching brain states, whereas ACh has the opposite effect by deepening these landscapes (Munn, Müller, Wainstein, & Shine, 2021). Overall, the LC and BF are synergistically connected and vital for cognitive flexibility.

Assessing these systems in a context where they naturally change provides an opportunity to observe variation in it. One such context is healthy aging, in which there are changes to attentional orienting (Madden & Langley, 2003), distractibility (Berti et al., 2013), and to the structural integrity of these neuromodulatory systems (Mather & Harley, 2016). The LC and BF are some of the first regions to show evidence of pathology in aging (Weinshenker, 2008; Zarow et al., 2003; Liu et al., 2015; Beardmore et al., 2021; Dahl et al., 2021) and this likely creates cognitive vulnerabilities years prior to the development of cognitive impairment, as has been demonstrated in Alzheimer’s Disease (Teipel et al., 2014; Kerbler et al., 2015; Jacobs et al., 2019; Dutt et al., 2020). Despite evidence of structural alterations in these neuromodulatory systems, it is unknown how these regions function within a network and how network structure changes with age to impact attentional processing. Advances in magnetic resonance imaging (MRI) allow us to investigate these small nuclei in humans. Turbo-spin echo (TSE) imaging provides a technique for structural localization of the small LC region (Keren et al., 2009; Turker et al., 2021) and multi-echo fMRI has been demonstrated to obtain high signal in the BF and LC (Markello et al., 2018; Turker et al., 2021). We use these tools to assess the agerelated changes in the interplay of the LC and BF nuclei and their regulation of the HPC and PCC during a go/no go task.

## Methods

### Participants

We examined 85 participants (36 younger adults, 14 middle-aged adults, 35 older adults) who completed this task as a part of a larger study that included neuropsychological assessment, structural and functional MRI scans, and several other cognitive tasks. Pupillary data and analyses from the same participants have been previously published in Riley et al. (2023). MRI data was not available for 18 participants because they did not complete all of the task, the acquired data was unusable due to technical problems, or their functional data did not converge during the ME-ICA pipeline (see Section 2.5).

### Participant Characteristics

Our ultimate dataset consisted of 30 younger (aged 19-45; average 25.16 years old; 64.5% female), 14 middle-aged adults (aged 46-65; average 58.14 years old; 61.2% female) and 23 older adults (aged 66-86; average 70.48 years old; 50% female) for a final sample of 67 healthy adults. Participants were screened for diagnosed cognitive impairment, neurological disease, head injury, ocular disease, and had vision and hearing that were normal or correctible-to-normal. Left-handed participants made up 6% of younger adults (2 participants), 14% of middle-aged adults (2 participants) and 8% of older adults (2 participants). Younger, middle-aged, and older adults had an average of 17.2y (SD = 3.1), 17.2y (SD = 3.1), and 17.5y (SD = 2.9) of education respectively. All participants were screened for cognitive impairment with the Montreal Cognitive Assessment. Younger adults had an average score of 27.9 (SD = 1.6, range 25-30, zero below cutoff), middle-aged adults had an average score of 27.2 (SD = 1.9, range 25-30, one below cutoff), and older adults had an average score of 27.0 (SD = 2.4, range 25-30, four below the cutoff after adjustment for years of education). None had a diagnosis of cognitive impairment of any kind. Participants were also given the Trail Making Test Part B, with an average time of 70.3s (SD = 23.7, range 41-119) for younger adults, 81.7s (SD = 37.8, range 39-177) for middle-aged adults and 95.4s (SD = 24.0, range 50-151) for older adults. While older adults had slower times to completion, none were longer than the predefined cutoff of 180 seconds.

Any participants that used vision correction were either given MR-safe lenses during testing, or, if they only used vision correction for reading, were given a brief vision test before entering the scanner to ensure that they would be able to see the task stimuli in focus.

### Task Overview

Detailed descriptions of the task and stimuli are presented in Riley et al. (2023), but are recounted here for completeness. Participants were asked to remember a series of pictures for a later test while performing a go/no-go auditory discrimination task. Participants listened for two types of tones (low and high) and responded by pressing a button for the target tone, but not the distractor tone. Participants completed 4 blocks of the task with the identity of the target switching each time. Detecting a target in the task has previously been shown to engage brain regions involved in attentional orienting, including the LC facilitating the processing of concurrently presented events relative to both distractor and baseline conditions. The identification of a target has been shown to increase the attentional and memory salience for concurrently presented events, as well as elicit activity in the LC relative to both distractors and no tone conditions (Swallow & Jiang, 2010; Swallow & Jiang, 2014; for a review, see Swallow et al., 2022).

### Task Stimuli

Tone stimuli were either high (1200 Hz) or low (400 Hz) and were 60 ms duration. Background visual stimuli were presented to maintain a consistent level of luminance and cognitive engagement across the testing session. They consisted of 144 color pictures and were evenly divided among pictures of faces, objects, and scenes. We generated an additional 144 scrambled image masks derived from the source images. The images were acquired from online resources (Huang, Jang, & Learned-Miller, 2007; http://vision.stanford.edu/projects/sceneclassification/resources.html) and personal collections. Between trials, the scrambled masks were presented to maintain light stimulation and were created by dividing an image into 256 squares and randomly shuffling them. Pixel intensities, both mean and variance, were matched across images using the SHINE toolbox (Willenbockel et al., 2010).

### Task Procedure

All participants performed the task as part of a longer MRI protocol. Each participant completed 4 blocks of 6 min 47 s each, for a total duration of less than 30 min, with brief breaks. On each 1.25 s long trial, one image (7 × 7 visual degrees; 256 × 256 pixels) was presented for 625 ms and immediately followed by a scrambled version of that same image for another 625 ms, and in some cases further scrambled images, also for 625ms. This timing, with no blank screen between trials, encouraged vigilance and rapid response times to help equate performance in younger and older participants.

On task trials (144 per block), participants first saw a picture and then a scrambled version of the same picture. Task trials were designated as target, distractor, or no tone trials in equal numbers. Participants were instructed that memory for the pictures would be tested later. Participants were asked to maintain fixation on a dot (0.25 visual degree diameter, red) at the center of the picture throughout the testing session. All 144 images were presented one time per block for a total of 4 repetitions across blocks and 576 total task trials. On task trials, either a high- or low-pitch or no tone played. Participants were told which was the target tone pitch, and this alternated across blocks, with the starting target tone counterbalanced across participants. When participants heard the specified target pitch for that run, participants pressed a button with their dominant hand pointer finger. Participants were instructed to make no motor response on trials with a distractor tone or no tone. Before the experiment, participants practiced the task. Tone volume was adjusted during a mock scan to ensure that participants were able to hear the tone over scanner noise. Tone sound level was always set to a standard to begin with and was raised only if participants were not able to discern the two different tones, with sound level ranging between 89% and 92% of maximum across participants.

From the perspective of the participant, there was a constant stream of scrambled images interspersed with intact pictures. Isoluminant changing and distinct background scrambled images, 164 per block without sound, were the majority of events to promote relatively constant low-level visual stimulation for pupil response measurement. These 164 scrambled images were in addition to the 144 pictures associated with trials and the 144 scrambled masks of each that followed it. The additional scrambled images also served to increase the unpredictability of the task trials. The median interval between true non-scrambled task trials was 2.5 s.

The go/no go task had a 3 × 6 design, with within-subject factors of tone type (no tone, distractor tone, target tone) and image type (female face, male face, beach, forest, car, chair); the latter included to examine potential image category effects. No tone trials were not examined in the analyses presented here. The trial sequence, specifically, the order and timing of each of the 18 trial types, was optimized using the AFNI function make_random_timing to produce sequences that maximized orthogonality of overlapping BOLD responses across trials and minimized the amount of unexplained variance in a simulated task. Inter-trial intervals were filled with scrambled images, as described above.

### MRI Acquisition and Preprocessing

Imaging was carried out at the Cornell University MRI Facility with a GE discovery MR750 3T scanner and a 32-channel head coil. Participants laid supine on the scanner bed with their head supported and stabilized. Ear plugs, headphones, and a microphone were used to reduce scanner noise, allow the participant to communicate with the experimenters, and to present auditory stimuli during the tasks. Visual stimuli were presented with a 32” Nordic Neuro Lab liquid crystal display (1920 × 1080 pixels, 60 Hz, 6.5 ms g to g) located at the back of the scanner bore and viewed through a mirror attached to the head coil. Pulse oximetry and respiration were recorded throughout all scans.

The imaging protocol consisted of a multi-echo acquisition (TR = 2500; TEs = 12.3, 26.0, and 40.0 ms; flip angle = 90°; matrix = 72 x 72; fov = 21 cm; slice thickness = 3.0 mm) and a structural T1-weighted MPRAGE (TR/TE = 7/3.42 ms; flip angle = 7°; matrix = 256 x 256; fov = 24 cm; slice thickness = 1 mm isotropic voxels). High resolution images of the LC were acquired with neuromelanin sensitive T1-weighted turbo-spin echo (TSE) structural scans (scan resolution = 512 x 512 mm; fov = 22.0 mm x 132.00 mm; TE = 11.26 ms; TR = 700 ms; flip angle = 120°). Visual stimuli were presented on a screen with a mirror, auditory stimuli were presented to participants via headphones, and participants responded to various tasks in the scanner by pressing a button box. Participants subsequently completed additional anatomical and functional scans that will be reported separately.

Preprocessing of the structural MRI images included normalization, skull stripping, segmentation, and spatial smoothing. Before preprocessing, EPI data was warped to MNI space. Processing of the multi-echo EPI data was performed using a preprocessing pipeline from AFNI (afni_procy.py; tedana.py, Version 2.4 beta 11) with the following blocks used: despike, tshift, align, tlrc, volreg, mask, combine, and scale (Taylor et al., 2018). Echo combination was completed using AFNI’s multi-echo ICA script (tedana.py, version 0.0.12). After preprocessing, the EPI data was spatially blurred (to 6 mm FWHM). This minimal amount of blurring has been shown to not critically change the ability to isolate LC activity (Turker et al., 2021). Quality control checks were completed at each stage in preprocessing to ensure accurate completion of each step.

### MRI Processing: Regions of Interest

LC ROIs were created for each participant from their neuromelanin sensitive TSE scan as described in Turker et al. (2021). The ROIs for the BF were taken from a probabilistic ROI obtained from a previous study (Zaborszky et al., 2008). Separate BFs for BF Ch1-3 (MS) and BF Ch4 (nbM) were obtained. Both BF ROIs were warped to native N27 space and thresholded. Left and right hippocampal and left and right posterior cingulate cortical (PCC) ROIs for each participant were obtained from FreeSurfer’s automatic parcellation. Volumes were reviewed for accuracy and masks were manually edited if necessary. After extraction, each participant’s hippocampal and PCC ROIs were thresholded to eliminate occasional nonzero voxels introduced during the warping process. Probabilistic hippocampal and PCC ROIs were obtained by finding the union set of all participants’ specific ROIs. ROIs in standard MNI space are shown in Figure 1. As hippocampal volume has shown to change with age, we ensured good overlap in hippocampal masks between each age group. The average hippocampal mask for each age group was formed and then the dice coefficient between all pairs of age groups was calculated to assess overlap between groups. All age group pairs showed good overlap in average hippocampal masks (young-middle: dice=0.751; young-old: dice=0.734; middle-old: dice=0.729).

**Figure 1.**
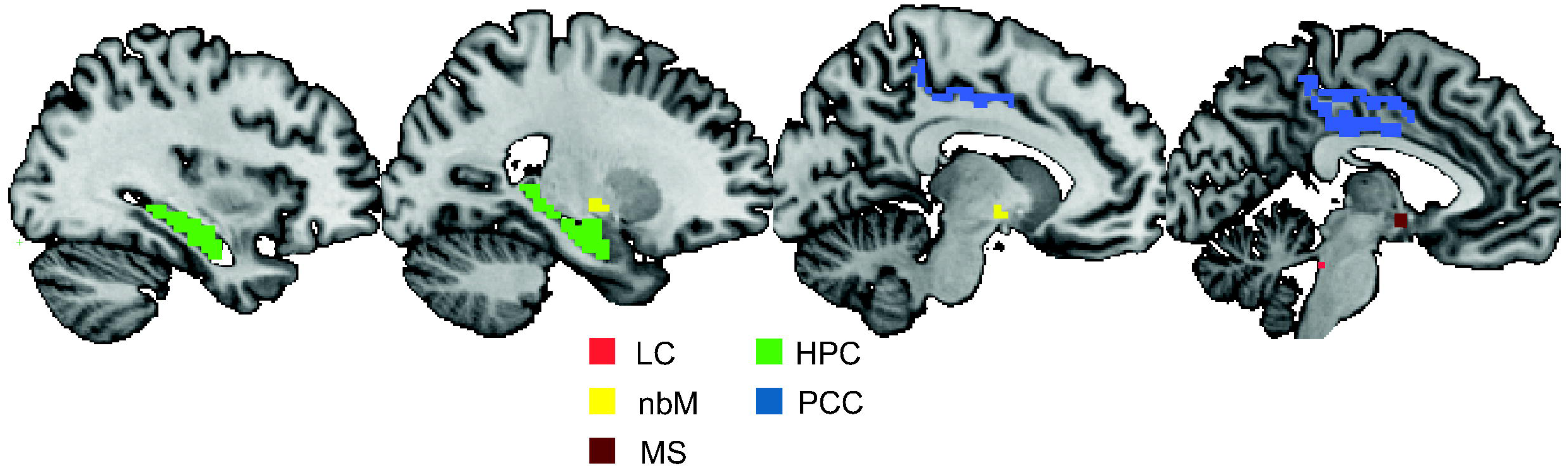
All ROIs in standard MNI space. HPC and PCC ROIs are segmented from FreeSurfer’s parcellation. nbM and MS ROIs are from a probabilistic atlas. LC ROIs are derived from neuromelanin-sensitive TSE scans. The HPC, PCC, nbM, MS, and LC ROIs above are from an example subject in standard MNI space. LC=locus coeruleus; HPC=hippocampus; nbM = Nucleus Basalis of Meynert; PCC=posterior cingulate cortex; MS=medial septum

### MRI Processing: Task-Dependent Functional Connectivity

To assess task-dependent functional connectivity between the LC, BF, and other target ROIs we conducted a generalized psychophysiological interaction (gPPI) analysis. This analysis was used because of the three task conditions and rapid event-related design of our go/no go task (McLaren et al., 2012; O’Reilly et al., 2012; Harrison et al., 2017). gPPI allowed us to characterize changes in LC, nbM, and MS functional coupling with other areas as they relate to the three go/no go task trial conditions (1:target, 2:distractor, 3:no tone). Briefly, the beta series for the LC ROI was extracted using a least-squares-sum (LSS) estimation approach (3dLSS in AFNI), generating one beta per trial. Next, for gPPI analysis, the beta series across each participant’s LC ROI was extracted (3dmaskave in AFNI). A canonical gamma hemodynamic response function (HRF) was convolved with the participant’s target tone stimulus timing file as input. Then the LC beta series was deconvolved with the stimulus-specific gamma HRF (waver in AFNI). A stimulus coding file was then generated that was the length of the number of trials in a single run, but each time point was 1 if that trial was a target tone or 0 if that trial was a distractor tone or no tone. The interaction between the deconvolved LC beta series and the stimulus coding file was then assessed by multiplying the two files (1deval -expr ‘a*b’ in AFNI). The interaction time series was then obtained by convolving the previous step’s output with a canonical gamma HRF. The above steps were repeated for each participant’s distractor trial time series separately, resulting in each participant having two separate interaction beta time series’. These interaction time series’ were then entered into a general linear model with the LC beta series (3dDeconvolve in AFNI) and interaction term beta weights were extracted for each trial type. Interaction term beta weights (referred to as “gPPI parameter estimates” going forward), for each trial type are interpreted as that specific trial’s task-dependent functional coupling for all voxels with the LC. The above steps were repeated with the nbM and MS ROIs as seeds to generate attention-dependent functional connectivity estimates across all three subcortical and brainstem neuromodulatory regions.

We operationally define target-related connectivity as task-dependent functional connectivity (gPPI parameter estimates) that is greater during target relative to distractor trials and we define distractor-related connectivity as connectivity that is greater during distractor relative to target trials. These definitions are useful for quantifying how trial-by-trial connectivity between neuromodulatory regions and its afferents support these two different aspects of the go/no-go task.

### Statistical Analysis: Dummy Coding

In analyses in which tone type (distractor or target) was a predictor of outcomes, distractors were coded as 1 and targets as 2. For age groups, younger adults were coded as 0, middle-aged adults as 1, and older adults as 2.

### Statistical Analysis Models

Several linear mixed effects models were completed in AFNI (3dLME) to assess gPPI parameter estimates in relation to several within- and between-subjects variables, while taking into account the random effect of subject. A linear mixed effects model was completed on the gPPI parameter estimates to assess changes in task-dependent functional coupling for each seed ROI in relation to trial type condition and age group. Cluster correction on the linear mixed effects results was completed to find the cluster size threshold (in voxels) and to find significant clusters (3dClusterize AFNI). Small-volume correction was completed using AFNI’s 3dClustSim to compute a threshold for a voxelwise p-value given the surviving cluster size. Note that our linear mixed effects results produce whole-brain voxelwise outputs, but we only report here results from our specific regions of interest.

## Results

### LC-seeded network

To determine how LC functional connectivity with its known afferents changed across conditions and age groups in our go/no go task, a linear mixed effects model was used. GPPI interaction parameter estimates for each trial, by tone type and age group, and including their interaction (tone x age group), were entered into the model.

### LC-BF nucleus basalis of Meynert (nbM) functional connectivity

Significant clusters in the nbM represented significant LC-nbM functional connectivity changes. There was a significant nbM cluster for the main effect of age group across all trial types (X,Y,Z = 28, −3, −12; cluster size = 16 voxels; F = 3.94; p = 0.035), but not for the main effect of trial type. There was a significant nbM cluster for the age-by-trial type interaction (X,Y,Z = −19, 8, −6; cluster size = 15 voxels; z = −2.33; p = 0.04), indicating a change in task-dependent LC-nbM coupling across age groups. The cluster that was significant for the age-by-trial type interaction did not overlap with the cluster significant for the main effect of age group, indicating two distinct nbM neuronal populations with coupling to the LC that changes across the lifespan.

To investigate the age-by-trial type interaction further, we extracted gPPI parameters from all subjects within the significant nbM cluster. We first computed one-sample t-tests within each age group to assess if task-dependent coupling within a given age group was stronger in targets or distractors. We found that in younger adults the LC-nbM connectivity was marginally greater during distractors (t = −1.813, p = 0.08), in middle-aged adults was significantly greater during distractors (t = −3.67, p = 0.0025), and in older adults reversed to being significantly greater during targets (t = 2.43, p = 0.022) (*Figure 2*). In comparing across age groups, LC-nbM coupling in older adults was significantly greater in targets versus distractors compared to younger and middle-aged adults (t = −3.13, p = 0.002; t = −3.44, p = 0.001). LC-nbM coupling was not significantly different between younger and middle-aged adults (t = 1.41, p = 0.165). The change in LC-nbM functional connectivity between targets and distractors had a marginally significant positive correlation with age (r = 0.228, p = 0.056). Task dependent LC-nbM coupling is ultimately greater during distractor trials in younger and middle-aged adults, but significantly changed in older adults.

**Figure 2.**
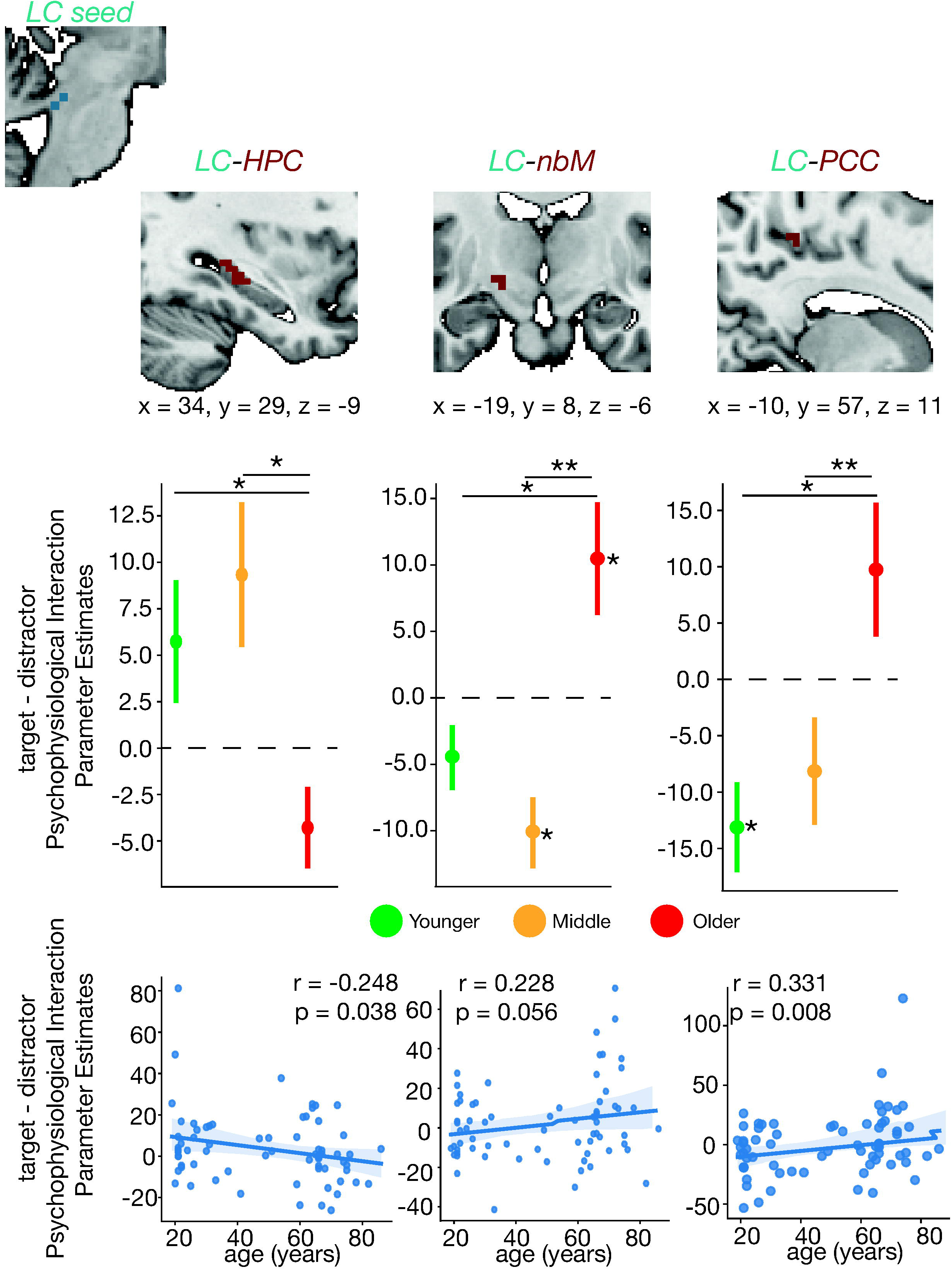
LC-seeded task-dependent functional connectivity across the lifespan. Generalized psychophysiological interaction (gPPI) parameter estimates were extracted from the voxels within significant ROIs (*refer to Methods Section: Task-Dependent Functional Connectivity*). Regions of interest with a significant age-by-trial type interaction for task-dependent functional connectivity with the LC are shown. Note this is not an exhaustive figure including all ROIs with significant age-by-trial type interactions (top row) Significant clusters from the age-by-trial type interaction. Clusters are displayed and coordinates are listed in MNI N27 space. (middle row) For each cluster, the difference between target and distractor trial psychophysiological interaction parameter estimates are plotted. Each column displays the results for a given region. An asterisk to the right of a single colored bar indicates a significant difference from zero (no task-dependent connectivity changes) with a one-sample t-test. Asterisks spanning two colored bars indicate a significant difference across two age groups computed with a two-sample t-test. All statistical tests were corrected for multiple comparisons using Bonferroni correction. (bottom row) Correlation between gPPI parameter estimates and age are also shown with the corresponding Pearson coefficient and p-value. LC-nbM functional connectivity had a significant age-by-trial type interaction such that the difference in connectivity between target and distractors was decreased in older adults compared to young and middle-aged adults. LC-HPC functional connectivity between target and distractor trials was greater in older adults compared to younger adults. LC-PCC functional connectivity was decreased in older adults compared to young and middle-aged adults. There was no significant LC-MS functional connectivity. LC=locus coeruleus; HPC=hippocampus; nbM = Nucleus Basalis of Meynert; PCC=posterior cingulate cortex; MS=medial septum *nbM-seeded network*

### LC-hippocampus (HPC) functional connectivity

Significant clusters in the HPC represent significant LC-HPC functional connectivity changes across age group or trial type. There were no significant clusters in the HPC for the main effect of age nor the main effect of trial type. However, there was a significant HPC cluster for the trial type-by-age interaction (X,Y,Z = 34, 29, −9; cluster size = 18 voxels; z = 2.96; p = 0.017).

Upon further investigation, LC-HPC in younger adults indicated marginal changes in connectivity across trial types, trending towards being greater during target trials (t = 1.71, p = 0.097). In middle aged adults, connectivity was significantly greater during target trials (t = 2.31, p = 0.036), and in older adults was marginally greater during distractor trials (t = −1.91, p = 0.067). LC-HPC coupling for targets compared to distractors was significantly weaker in older adults compared to younger (t = 2.38, p = 0.020) and middle-aged adults (t = 3.21, p = 0.002), with no significant difference between younger and middle-aged adults (t = −0.65, p = 0.518) (*Figure 2*). The change in LC-HPC functional connectivity between targets and distractors had a significant negative linear correlation with age (r = −0.248, p = 0.038). Task-dependent LC-HPC coupling was marginally greater during target trials in young and middle-aged adults, but significantly decreased in older adults to marginally support distractor processing instead.

### LC-Posterior Cingulate Cortex (PCC)

Significant clusters in the PCC represent significant LC-PCC functional connectivity changes. There were no significant clusters in the PCC for the main effect of age group nor for the main effect of trial type. There was a significant PCC cluster for the trial type-by-age interaction (X,Y,Z = −10, 57, 11; cluster size = 46 voxels; z = −2.41; p = 0.036).

One-sample t-tests revealed that in younger adults LC-PCC coupling was greater during distractor trials (t = −3.23, p = 0.003), whereas LC-PCC coupling in middle-aged and older adults was not significantly different across the two trial types (t = −1.657, p = 0.119; t = 1.602, p = 0.112). LC-PCC coupling for targets compared to distractors was significantly greater in older adults compared to younger adults (t = −3.217, p = 0.002) and compared to middle-aged adults (t = −3.446, p = 0.0014), with no reliable difference between younger and middle-aged adults (t = −0.74, p = 0.463) (*Figure 2*). The change in LC-PCC functional connectivity between targets and distractors had a significant positive linear correlation with age (r = 0.331, p = 0.008). The results overall indicate that LC-PCC coupling was greater during distractor trials in younger adults, but no longer task-dependent by old age.

There was no significant LC-MS (BF Ch1-3) coupling for any main effects nor interactions. This aligns with previous literature indicating structural connections between the LC and nbM (Espana & Berridge, 2006), but not between the LC and MS. Overall, with the LC as a seed region, gPPI analysis revealed that across age LC coupling with the nbM, HPC, and PCC changed in a task-dependent manner. Namely, with age LC-nbM and LC-PCC functional coupling was greater during distractor trials in young and middle age but reversed in old age to support target trials. On the other hand, LC-HPC functional coupling was greater during target trials in young and middle age, but reversed to support distractor trials in old age. There are clear task-dependent changes in LC coupling with both subcortical and cortical regions that markedly change with age most strikingly later in life.

### nBM-seeded network

To determine how nbM functional connectivity with its known afferents changed across conditions and age groups in our go/no go task, a linear mixed effects model was used. GPPI parameter estimates for each trial, by tone type and age group, and including their interaction (tone x age group), were entered into the model.

#### nBM-LC

Significant clusters in the LC represent significant nbM-LC functional connectivity changes. There were no significant LC clusters for the main effect of age group nor trial type, but there was a significant LC cluster for the trial type-by-age interaction (X,Y,Z = −4, 34, −27; cluster size = 22 voxels; z = 2.776; p = 0.034).

One-sample t-tests revealed that in younger adults, LC-nbM connectivity did not depend on trial type (t = 1.369, p = 0.181) (*Figure 3*). However, in middle-aged adults, nbM-LC coupling was stronger during targets (t = 2.181, p = 0.0466). In older adults nbM- LC coupling reversed and was actually stronger during distractor trials (t = −3.694, p = 0.0012). nbM-LC coupling between targets and distractors was greater in younger and middle-aged adults compared to older adults (t = 3.495, p = 0.0009; t = 3.827, p = 0.0004). Additionally, middle-aged adults had marginally greater nbM-LC coupling compared to younger adults (t = −1.944, p = 0.058). The change in LC-nbM functional connectivity between targets and distractors did not have a significant linear correlation with age (r = −0.138, p = 0.255). However, when testing for any quadratic effects there was a significant curvilinear relationship between nbM-HPC functional connectivity with age (p = 0.005). Task-dependent nbM-LC coupling, with nbM as the seed, was stronger during target trials in middle age but strongly reversed in old age to be greater during distractor trials. Of note, we observe differences in nbM-LC coupling between all pairs of the three age groups, indicating strong sensitivity of task-dependent nbM-LC coupling to age.

**Figure 3.**
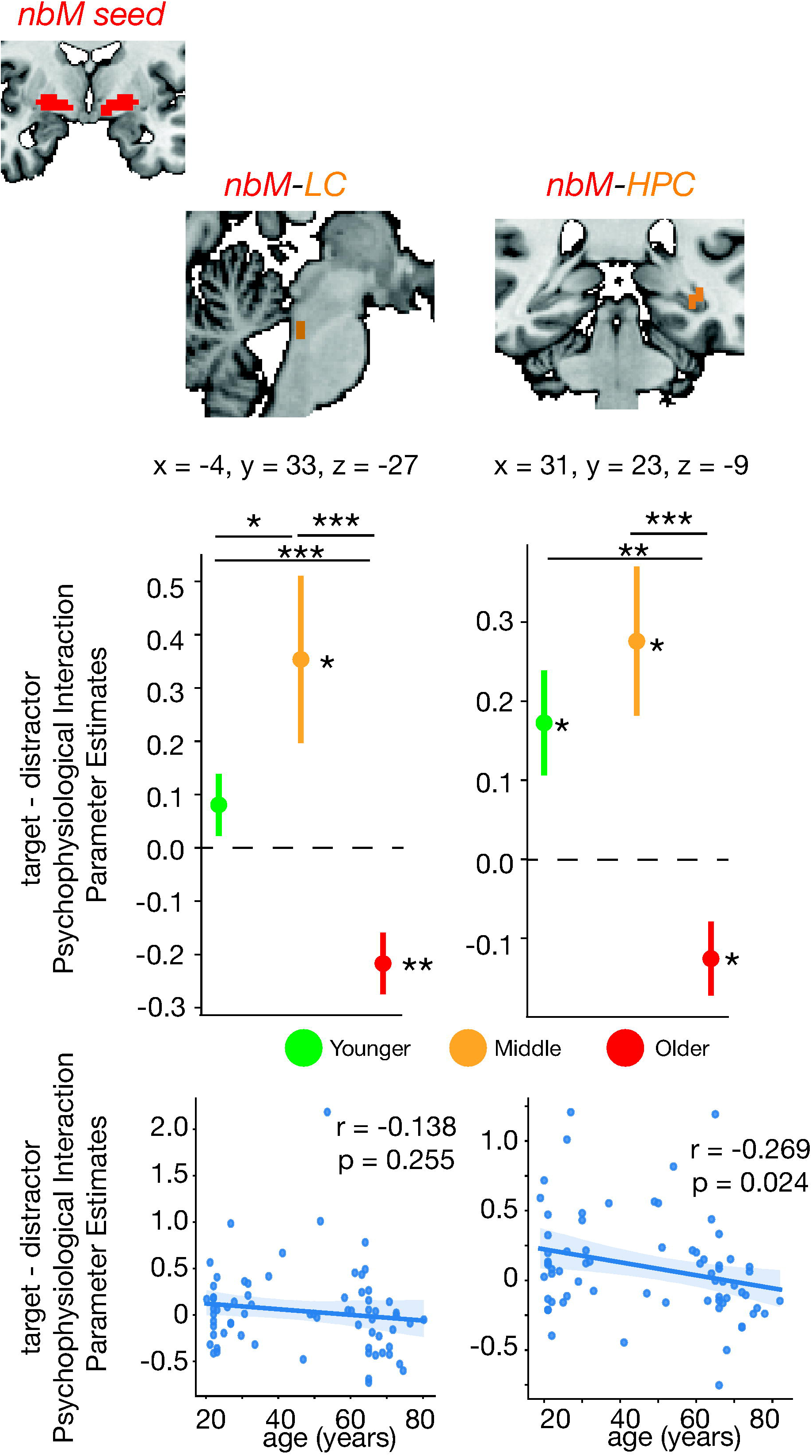
nbM-seeded task-dependent functional connectivity across the lifespan. Generalized psychophysiological interaction (gPPI) parameter estimates were extracted from the voxels within significant ROIs (*refer to Methods Section: Task-Dependent Functional Connectivity*). Regions with a significant age-by-trial type interaction for task-dependent functional connectivity with the nbM are shown. (top row) Significant clusters from the age-by-trial type interaction. Clusters are displayed and coordinates are listed in MNI space. (middle row) For each cluster, the difference between target and distractor trial gPPI estimates are plotted. Each column displays the results for a given region. An asterisk to the right of a single colored bar indicates a significant difference from zero with a one-sample t-test. Asterisks spanning two colored bars indicate a significant difference across two age groups computed with a two-sample t-test. All statistical tests were corrected for multiple comparisons using Bonferroni correction. (bottom row) The correlation of the parameter estimates and age are also shown with the corresponding Pearson coefficient and p-value. nbM-LC and nbM-HPC functional connectivity had a significant age-by-trial type interaction such that the difference in connectivity between target and distractors was decreased in older adults compared to young and middle-aged adults. There was no significant LC-MS functional connectivity nor nbM-PCC functional connectivity. LC=locus coeruleus; HPC=hippocampus; nbM = Nucleus Basalis of Meynert; PCC=posterior cingulate cortex; MS=medial septum

#### nBM-HPC

Significant clusters in the HPC represented significant nbM-HPC functional connectivity changes. There were significant bilateral HPC clusters for the main effect of trial type group across all age groups (*left HPC*: X,Y,Z = 43, 26, −12; cluster size = 20 voxels; F = 5.901; p = 0.0142; *right HPC*: X,Y,Z = −31, 23, −9; cluster size = 29 voxels; F = 7.214; p = 0.009), suggesting that nbM coupling with the left and right HPC is related to rapid changes according to trial type. Additionally, there was a significant HPC cluster for the main effect of age group (X,Y,Z = 23, 32, −13; cluster size = 38 voxels; F = 3.54; p = 0.045), indicating that task-dependent nbM-HPC coupling changes across the lifespan. Finally, there was a significant HPC cluster in the left hemisphere for the trial type-by-age interaction (X,Y,Z = 31, 23, −9; cluster size = 95 voxels; z = 3.153; p = 0.0153).

Upon further investigation, nbM-HPC coupling was significantly greater during target trials for younger and middle-aged adults (t = 2.56, p = 0.0157; t = 2.824, p = 0.0135). However, nbM-HPC coupling was significantly greater during distractor trials for older adults (t = −2.635, p = 0.0151) (*Figure 3*). Across age groups, nbM-left HPC coupling between targets and distractors in older adults significantly decreased compared to younger and middle-aged adults (t = 3.37, p = 0.0014; t = 4.086, p = 0.0002). There was no significant difference in nbM-HPC coupling between young and middle-aged adults (t = −0.872, p = 0.387). The change in nbM-HPC functional connectivity between targets and distractors had a significant negative correlation with years of age (r = −0.269, p = 0.024). When testing for any quadratic effects there was a significant curvilinear relationship between nbM-HPC functional connectivity with age (p = 0.0458), but this does not survive multiple comparisons correction. Similar to nbM-LC coupling, task-dependent nbM-HPC coupling was greater during target trials in younger and middle-aged adults, but markedly reversed to be greater during distractor trials in older age.

There was no significant nbM-MS task-dependent coupling for any main effects nor interactions. This aligns with the lack of previous literature indicating structural connections between the nbM and MS. There was also no significant nbM-PCC coupling for any main effects nor interactions. Overall, with the nbM as a seed region, gPPI analysis revealed that nbM coupling with the LC and bilaterally with the HPC interacts with trial type and age. Further examination reveals similar age differences in nbM-LC and nbM-HPC coupling, with increased connectivity during target trials until old age in which there is increased connectivity during distractor trials. There are clear task-dependent changes in nbM coupling with the LC and HPC that markedly change with age, especially later in life.

### MS-seeded network

GPPI interaction parameter estimates for trials, by tone type and age group and their interaction (tone x age group), during the go/no go task were entered into a linear mixed effects model to assess MS task-dependent functional connectivity with known afferents.

#### MS-HPC

Significant clusters in the HPC represented MS-HPC functional connectivity changes across task conditions. There was a significant HPC cluster for the main effect of age group across all trial types (X,Y,Z = −31, 6, −28; cluster size = 21 voxels; F = 7.455; p = 0.035) and a cluster for the main effect of trial type (X,Y,Z = 20, 27, −19; cluster size = 75 voxels; F = 6.675, p = 0.0217). There was a significant HPC cluster in the left hemisphere for the age-by-trial type interaction (X,Y,Z = 22, 23, −9; cluster size = 63 voxels; z = 2.905; p = 0.0062).

One-sample t-tests revealed that in young and middle-aged adults, MS-HPC coupling was greater during targets relative to distractor trials (t = 2.384, p = 0.023; t = 3.370, p = 0.0045, respectively). In older adults, there was no difference between target and distractor MS-HPC connectivity (t = −0.943, p = 0.354). MS-HPC coupling differences between targets and distractors in older adults was significantly decreased compared to younger and middle-aged adults (t = 2.296, p = 0.025; t = 3.678, p = 0.0007, respectively). Additionally, middle-aged adults had greater MS-HPC coupling than younger adults (t = −2.145, p = 0.037) (*Figure 4*). The change in MS-HPC functional connectivity between targets and distractors did not have a significant linear correlation with age (r = −0.134, p = 0.264). The results overall suggest that task-dependent MS-HPC coupling is greater during target trials in younger adults, increases more in middle-aged adults, but in older adults is no longer different between target and distractor trials. Notably, MS-HPC coupling is significantly different across all age groups, suggesting that MS-HPC coupling dynamically changes across the entire lifespan, not just in old age.

**Figure 4.**
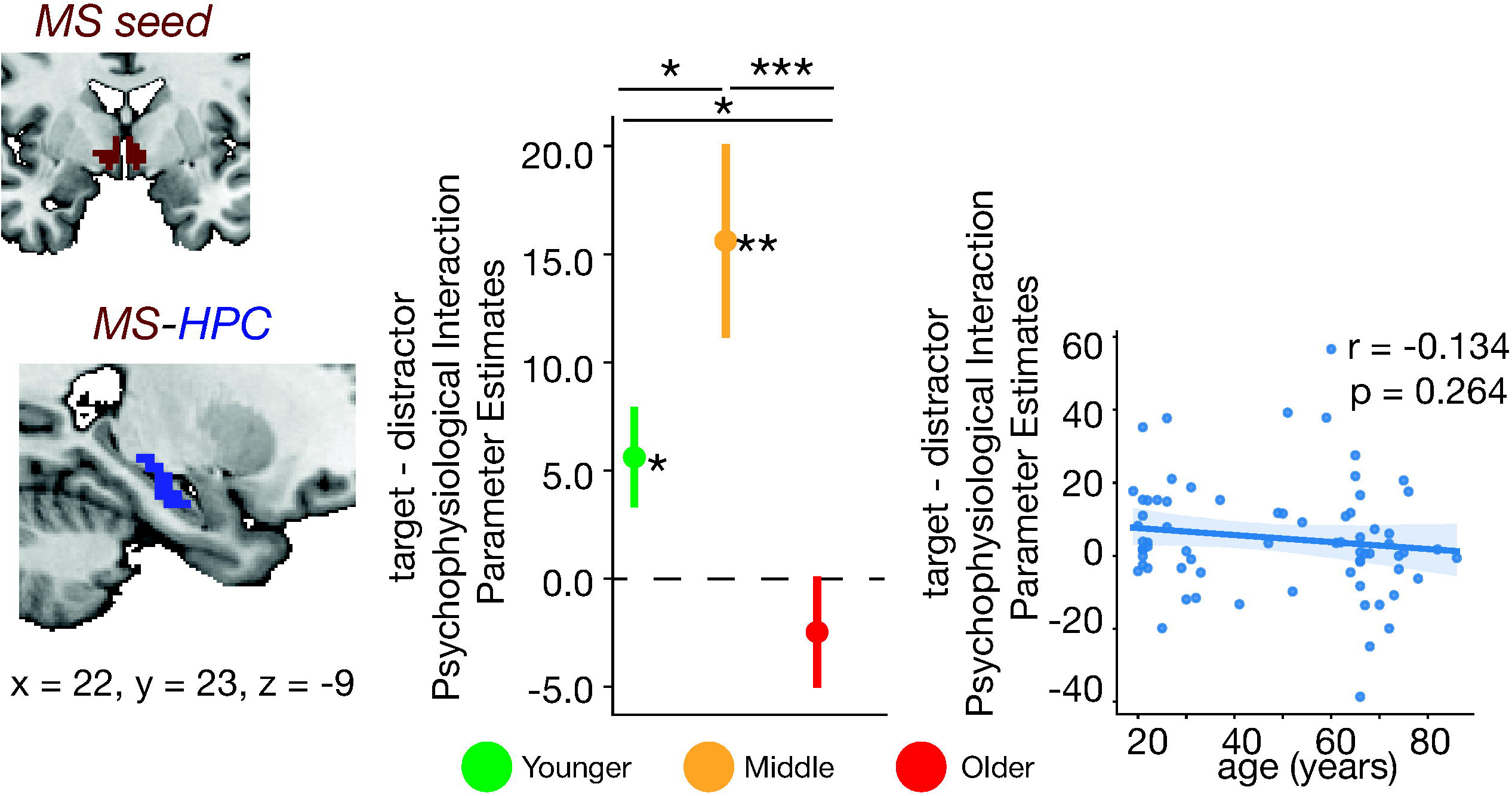
MS-seeded task-dependent functional connectivity across the lifespan. Generalized psychophysiological interaction (gPPI) parameter estimates were extracted from the voxels within significant ROIs (*refer to Methods Section: Task-Dependent Functional Connectivity*). Regions with a significant age-by-trial type interaction for task-dependent functional connectivity with the MS are shown. (top row) Significant clusters from the age-by-trial type interaction. Clusters are displayed and coordinates are listed in MNI N27 space. (middle row) For each cluster, the difference between target and distractor trial psychophysiological interaction parameter estimates are plotted. Each column displays the results for a given region. An asterisk to the right of a single colored bar indicates a significant difference from zero with a one-sample t-test. Asterisks spanning two colored bars indicate a significant difference across two age groups computed with a two-sample t-test. All statistical tests were corrected for multiple comparisons using Bonferroni correction. (bottom row) The correlation of the parameter estimates and age are also shown with the corresponding Pearson coefficient and p-value. MS-HPC functional connectivity had a significant age-by-trial type interaction such that the difference in connectivity between target and distractors increased from young to middle age, but decreased in old age. There was no significant MS-LC, MS-nbM, nor MS-PCC functional connectivity. LC=locus coeruleus; HPC=hippocampus; nbM = Nucleus Basalis of Meynert; PCC=posterior cingulate cortex; MS=medial septum

In line with the known structural connectivity of the MS, there were no significant MS-LC (Espana & Berridge, 2006), MS-nbM, nor MS-PCC task-dependent coupling for the main effect of age and trial type, nor the interaction of age and trial type. Altogether, MS-HPC task-dependent coupling is greater during target trials in younger adults, increases even more so in middle-aged adults, but markedly decreases in older adults where MS-HPC coupling is no longer differentiated between target and distractor trials. Thus, MS-HPC coupling changes across the lifespan with significant age differences present already in middle age.

### Summary of results

In younger adults, we observed significant task-related functional coupling in all seed regions, with the majority of task-dependent connections stronger in target (salient) trials. This network structure reconfigures in middle age, with functional connectivity increasing between many regions, with an overall increase in connections stronger in target trials (Figure 5). In middle-aged adults the strength of connectivity increases across several network edges, such as between the MS and HPC, whereas LC-PCC coupling is no longer significant. Further, in middle age LC-nbM coupling begins to encode a distracting signal. In old age, the network became much sparser, with only two node pairs with significant task-dependent connectivity, the nbM-HPC and LC-nbM. With old age nbM-HPC connectivity now encodes a distracting signal, whereas LC-nbM connectivity now encodes a salience signal, a complete reversal from middle age. Additionally, the ratio of target-related connections compared to distractor-related connections is most strongly reduced in older adults.

**Figure 5.**
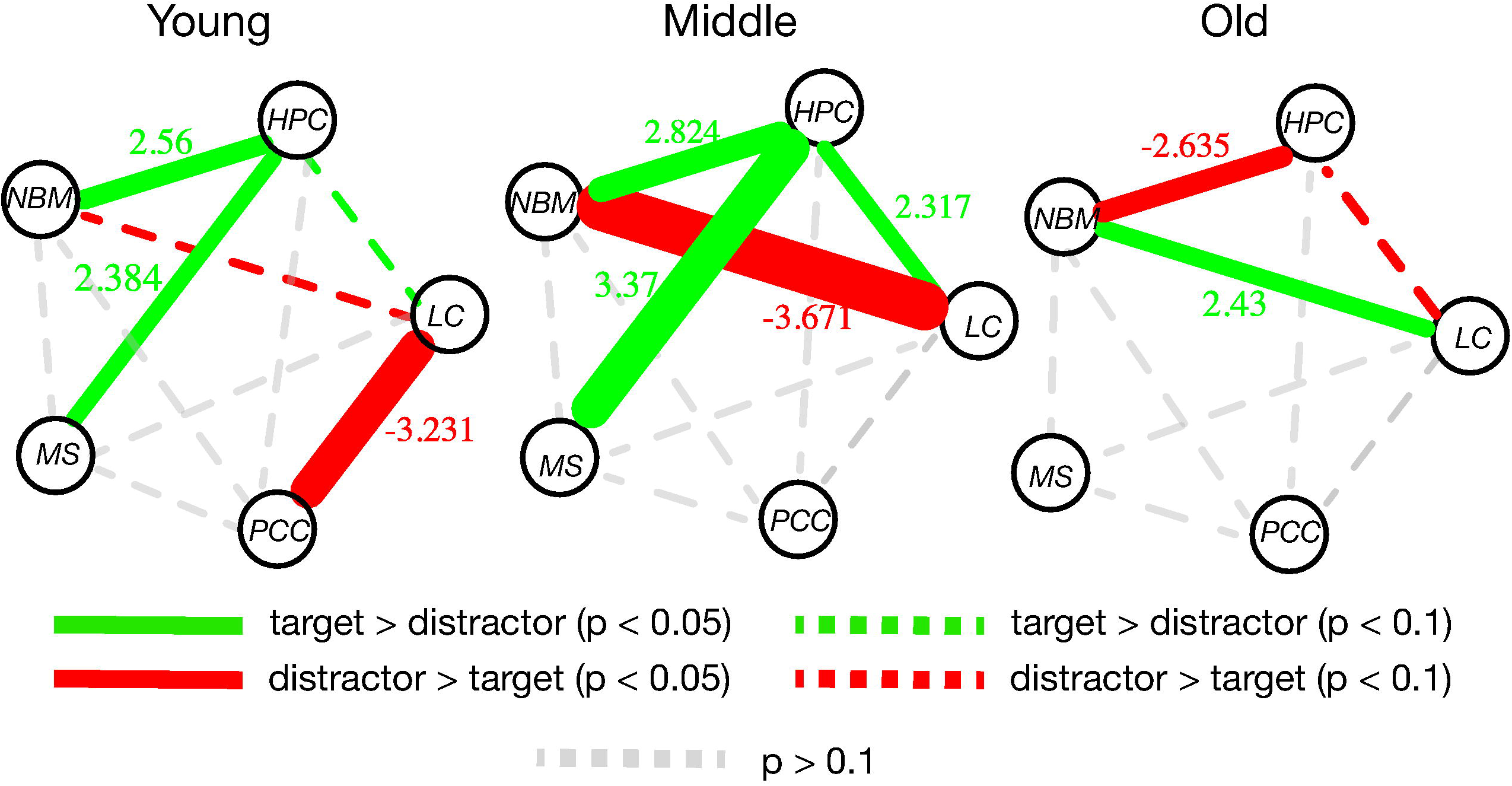
Path diagram summarizing the results from. Figures 2-4. Green lines indicate a significant relationship between functional coupling and task, with greater coupling during targets compared to distractors. For colored lines, line thickness and the values along the lines indicate the strength of the relationship, as measured by the t-statistic of the task-dependent coupling in each group calculated from a one-sample t-test against a mean of zero. Green lines indicate greater coupling during targets compared to distractors. Red lines indicate greater coupling during distractors compared to targets. Grey dotted lines indicate a relationship between functional coupling and task that had a p-value > 0.1. Color dotted lines indicate a relationship between functional coupling and task that had a p-value > 0.05 and p-value < 0.1. LC: locus coeruleus; nbM: nucleus basalis of Meynert; MS: medial septum; HPC: hippocampus; PCC: posterior cingulate cortex.

## Discussion

In this study we examined how age alters functional coupling between known afferent projection sites of noradrenergic and cholinergic subcortical nuclei during attentional orienting. We demonstrated changes in task-dependent functional connectivity amongst neuromodulatory regions which relate to attentional orienting and task switching on the timescale of seconds (Berridge & Waterhouse, 2003; Aston-Jones & Cohen, 2005). We found that functional connectivity between the nbM and HPC was greater during target trials than distractor trials in young and middle-aged adults, but in old age the reverse was true. In contrast, LC connectivity with the PCC and MS connectivity with the HPC was strongest during distractor trials and target trials, respectively, in younger individuals. LC connectivity to PCC and MS were no longer related to task condition by old age, indicating an age-related reduction in task-related connectivity across these regions. Altogether, task-dependent connectivity across this network of regions changed in both middle and old age, with an overall sparser and refocused network dominated by functional coupling between neuromodulatory nuclei in older adults. As successful attentional orienting requires complimentary salience and distractor signals to orient attention within a rapidly changing environment, with age the network supporting the push and pull between these signals is sparser. In old age this network appears to contain fewer nodes, resulting in an altered contribution of attentional gain relative to attentional tuning.

We restricted our analysis to nodes with known structural connectivity and we found functional coupling between the LC, BF, HPC, and PCC (Levitt & Moore, 1978; Mesulam et al., 1983; Espana & Berridge, 2006; Walling et al, 2011; Hagena, Hansen & Manahan-Vaughan, 2016; Hansen et al., 2017; Wagatsuma et al., 2017; Bacon, Pickering, & Mellor, 2020; James et al. 2020). As expected, these results indicate that attentional orienting coactivates regions with synaptic connections (Kellerman et al., 2012). The observed coupling also replicates previous findings of the resting-state functional connectivity between the LC and PCC (Jacobs et al., 2018; Turker et al., 2021), LC and HPC (Turker et al., 2021), LC and nbM (Jacobs et al., 2018), and nbM and PCC (Markello et al., 2018). Our finding that LC-HPC functional connectivity is numerically greater for target compared to distractor trials is in the same direction as a prior result using the same task (Moyal et al., 2022). Unlike previous studies which assessed age-related functional connectivity amongst this network during rest, we demonstrate functional coupling within this network that supports specific aspects of trial-by-trial changes in attentional state and is modulated by noradrenergic and cholinergic coactivation with several regions. In doing so, we show a pronounced age-related network reconfiguration which follows the known anatomical connections across these neuromodulatory regions.

Despite the age-related alterations in this network, the older adults in this study were all healthy with no diagnosed cognitive deficits. Although much evidence indicates cognitive decline with aging, not all aspects of cognition diminish with age and some domains may even improve with aging. Meta-analyses of psychophysiological and cognitive assessments performed in older adults indicate mixed results across decades of studies (Madden, 2007; Verissimo et al., 2022). Reaction time, distraction detection, and other attentional orienting performance metrics appear to diminish with old age, but other aspects of attention such as attentional control and decreased mind-wandering have been shown to be preserved through at least some stages of old age (Fountain-Zaragoza et al., 2018; Verissimo et al., 2022). These inconsistencies may be a result of large variability in the older population. Amongst potential mediators of this heterogeneity are educational experience and socioeconomic status (Verissimo et al. 2022). Though we cannot directly relate individual variability of our older sample to the age-related connectivity changes in these neuromodulatory networks, our results suggest that regions responsible for attentional orienting dynamically change across the lifespan. Further work is needed to characterize the relationship between heterogeneous cognitive outcomes in old age with functional and structural changes of these regions.

Although aspects of attention may be preserved with old age, much evidence indicates robust age-related deficits (Coubard et al., 2011; Cashdollar et al., 2013; Bier et al., 2017). We demonstrate a drastically sparser network structure in older adults which may account for some of these age-related deficits. For example, the sparser connectivity and weaker functional coupling supporting distractor signaling aligns with the finding that older adults have prolonged processing of distractors and weakened distractor detection (Cashdollar et al., 2013). Our results indicate that older adults have strong LC-nbM coupling supporting salience signals but a weak nbM-HPC coupling supporting distractor signals compared to distraction-supporting connectivity earlier in life, suggesting that this reconfigured attentional network may be the origin of stronger attentional gain without tuning (Riley et al., 2023). Although the network does become sparser with age, consistent with a monotonic decline, the functional connections between the LC and nbM becomes strengthened. As the NE and ACh systems have been proposed to synergistically support attentional switching and uncertainty in the environment, it holds that healthy older adults with sparse cortical connectivity with the LC and BF still have significant LC-BF functional coupling (Yu & Dayan, 2005; Munn, Müller, Wainstein, & Shine, 2021). In young and middle age, the LC and BF influence the attentional system through subcortical-cortical task-dependent functional connections. This diverse network architecture allows for the LC and BF to influence its afferents, namely the HPC and PCC, in a task-dependent manner, which has been previously shown to support faithful attentional and memory processing (Levitt & Moore, 1978; Mesulam et al., 1983; Detari, Sembra, & Rasmusson, 1997; Espana & Berridge, 2006; Markello et al., 2018). In the healthy older adults, we observed that neuromodulatory coupling with cortical afferents was largely reduced, with the LC and MS no longer having significant attention-dependent coupling with the HPC nor with the PCC, respectively. Since no attentional deficits were detected in this healthy older adult sample and LC-nbM functional coupling was one of the two significant network edges remaining in this age group, it is evident that these neuromodulatory nuclei more sparsely connect with their cortical afferents with age but maintain strong internuclei connectivity.

Despite being a potentially compensatory effect, this altered network architecture likely creates a vulnerable system in which cognitive abilities, such as attentional reorienting and task switching, may be at risk for becoming deficient (Peters, A., Setharas, C., & Luebke, J. I., 2008; Arnsten, Wang, & Paspalas, 2012). As mentioned previously, the LC and BF are some of the first regions to show evidence of structural and functional alterations with Alzheimer’s Disease (Zarow et al., 2003; Weinshenker, 2008; Teipel et al., 2014; Kerbler et al., 2015; Liu et al., 2015; Jacobs et al., 2019; Dutt et al., 2020; Beardmore et al., 2021; Dahl et al., 2021). Our study builds on a body of literature that overall demonstrates significant changes in these neuromodulatory systems in both healthy and pathological aging. Although our healthy older adult sample did not include any individuals with a diagnosis of cognitive impairment or serious neurological disease, there were likely many cases of prodromal neurodegenerative disease since roughly 50% of individuals who live into their 90s will be diagnosed with Alzheimer’s Disease (Gilsanz et al., 2019). Once symptoms of Alzheimer’s Disease reach a clinical level, damage to these neuromodulatory regions is already substantial (Braak, 2011) and treatment has limited effects on symptoms and time course of the disease. Given that the LC and BF are critical for cognitive impairment with age, investigating the functional coupling of the network we measured in older adults with diagnosed Mild Cognitive Impairment (MCI) and Alzheimer’s Disease (AD) will inform how disease onset and progression relates to restructuring of the LC and BF network along the same timescale. Investigating this functional network in older adults with cognitive impairments is a vital target for future work.

Our results additionally inform hypotheses about the synergistic role that the LC and nbM perform together to alter large-scale brain states. The balancing act that our brain upholds throughout our everyday life requires rapidly allocating attentional resources to the many inputs our system receives, including both external sensory and internally generated inputs. As it has been posited that neuromodulatory systems are at this interface of generating flexible brain states, it follows that these two systems require functional coupling to allow for rapid fluctuations in attention and downstream cognition (Shine, 2019; Munn et al., 2021). There is evidence to suggest that the balance between integration and segregation is related to white matter connectivity and that large-scale network shifts from phasic LC and nbM firing covary with the strength of the structural connectivity between the LC and nbM (Taylor et al., 2022). In line with this evidence, our results indicate strong LC-nbM functional connectivity during second-to-second changes in attentional orienting.

Several limitations to the current work can be further addressed in future work. First, our dataset is undersampled within the 46-65 years old middle age group and has high variance. As many structural and functional deficits can already be seen in middle age, investigating this age group is of great importance for interventional and preventative measures against age-related cognitive impairment (Jacobs et al., 2018). Future work will assess how attention-dependent functional connectivity between the LC and nbM changes with cognitive deficits, such as in MCI and AD, and how this coupling relates to disordered attentional control. As several studies have shown that functional connectivity measured with fMRI is strongly correlated with the underlying structural connectome (Honey et al., 2009; O’Reilly et al., 2013), we would expect that major structural changes in MCI and AD would subsequently interfere with this network’s task-dependent functional coupling and drastically perturb the network architecture we observed.

In conclusion, we find that the major sources of norepinephrine and acetylcholine of the brain, the LC and BF nuclei, and their known structural connections coordinate to support trial-by-trial attentional orienting and do so differently across the lifespan. In old age, these neuromodulatory networks supporting attentional regulation of afferent cortical regions become sparser; however, the mutual connectivity between neuromodulatory nuclei strengthens. Our findings provide new insights into how neuromodulatory nuclei support attention-state dependent functional coupling, and how their changes support attentional orienting with healthy aging.

## Code Accessibility

Customized software created for the gPPI analyses can be made available upon request.

## Acknowledgements

This work was supported by F32 AG058479 to ER and R01AG066430 to EDR and AKA. This research was carried out at the Cornell University Magnetic Resonance Imaging Facility within the Cornell Human Neuroscience Institute. Our experimental protocol was reviewed by the Cornell Institutional Review Board, protocol #1910009087. We thank Elizabeth Sharp, Love Nemecek, Julio Salas, and Elena Cabrera for assistance with data collection.

## References

Arnsten, A. F. T., Wang, M. J., & Paspalas, C. D. (2012). Neuromodulation of thought: Flexilities and vulnerabilities in prefrontal cortical network synapses. Neuron, 76, 233–239. doi: 10.1016/j.neuron.2012.08.038

Aston-Jones, G., & Cohen, J. D. (2005). An integrative theory of locus coeruleus-norepinephrine function: Adaptive gain and optimal performance. Annual Review of Neuroscience, 28(1), 403–450. 10.1146/annurev.neuro.28.061604.135709

Bacon, T., Pickering, A., & Mellor J. (2020). Noradrenaline release from locus coeruleus terminals in the hippocampus enhances excitation-spike coupling in CA1 pyramidal neurons via ɑ-adrenoreceptors. Cerebral Cortex, 30, 6135–6151. doi:10/1093/cercor/bhaa159

Beardmore, R., Hou, R., Darekar, A., Holmes, C., & Boche, D. (2021). The locus coeruleus in aging and Alzheimer’s Disease: A postmortem and brain imaging review. J Alzheimers Dis., 83, 5–22. doi: 10.3233/JAD-210191

Berridge, C. W. & Waterhouse, B. D. (2003). The locus coeruleus-noradrenergic system: modulation of behavioral state and state-dependent cognitive processes. Brain Research Reviews, 42, 33–84. doi: 10.1016/S0165-0173(03)00143-7

Berti, S., Grunwald, M., & Schroger, E. (2013). Age dependent changes of distractibility and reorienting of attention revisited: an event-related potential study. Brain Res., 1491, 156–166. doi: 10/1016/j.brainres.2012.11.009

Bier, B., Lecavalier, N. C., Malenfant, D., Peretz, I., & Bellevile, S. (2017). Effect of aging on attentional control in dual-tasking. Exp Aging Res., 43, 161–177. doi:10.1080/0361073X.2017.1276377

Cashdollar, N., Fukuda, K., Bocklage, A., Aurtenetxe, S., Vogel, E. K., & Gazzaley, A. (2013). Prolonged disengagement from attentional capture in normal aging. *Pyschol*. Aging, 28, 77–86. doi:10.1037/a0029899.

Chun, M. M. & Turk-Browne, N. B. (2007). Interactions between attention and memory. Current Opinion in Neurobiology, 17, 177–184. doi: 10.1016/j.conb.2007.03.005

Corbetta, M., Patel, G., & Shulman, G. L. (2008). The reorienting system of the human brain: From environment to theory of mind. Neuron, 58, P306–324. doi: 10.1016/j.neuron.2008.04.017

Coubard, O. A., Ferrufino, L., Boura, M., Gripon, A., Renaud, M., & Bherer, L. (2011). Attentional control in normal aging and Alzheimer’s disease. Neuropsychology, 25, 353–356. doi:10.1037/a0022058.

Dahl M. J., Mather, M., Düzel, S., Bodammer, N. C., Lindenberger. U., Kühn, S., Werkle-Bergner, M. (2019). Rostral locus coeruleus integrity is associated with better memory performance in older adults. Nat Hum Behav, 3, 1203–1214. PMID: 31501542

Detari, L., Semba, K., & Rasmusson, D. D. (1997). Responses of cortical EEG-related basal forebrain neurons to brainstem and sensory stimulation in urethane-anesthetized rats. European Journal of Neuroscience, 9, 1153–1161.

Dutt, S., Li, Y., Mather, M., & Nation, D. A. (2020). Brainstem volumetric integrity in preclinical and prodromal Alzheimer’s disease. J Alzheimers Dis., 77, 1579–1594. doi: 10.3233/JAD-200187

Espana, R., & Berridge, C. W. (2006). Organization of noradrenergic efferents to arousal-related basal forebrain structures. The Journal of Comparative Neurology, 496, 668–683. doi: 10.1002/cne.20946

Fountain-Zaragoza, S., Puccetti, N. A., Whitmoyer, P., & Prakash, R. S. (2018). Aging and attentional control: Examining the roles of mind-wandering propensity and dispositional mindfulness. J Int Neuropsychol Soc, 24, 876–888. doi:10.1017/S1355617718000553.

Gilzanz, P., Corrada, M. M., Kawas, C. H., Mayeda, E. R., Glymour, M. M., Quesenberry Jr., C. P., Lee, C., & Whitmer, R. A. (2019). Incidence of dementia after age 90 in a multiracial cohort. Alzheimer’s Dement., 15, 497–505. doi: 10.1016/j.jalz.2018.12.006

Hagena, H., Hansen, N., Managan-Vaughan, D. (2016). â-adrenergic control of hippocampal function: Subserving the choreography of synaptic information storage and memory. Cerebral Cortex, 26, 1349–1364. doi:10.1093/cercor/bhv30

Hansen, N. (2017). The longevity of hippocampus-dependent memory is orchestrated by the locus coeruleus-noradrenergic system. Neural Plasticity. doi:10.1155/2017/2727602

Honey, C. J., Sporns, O., Cammoun, L., Gigandet, X., Thiran, J. P., Meuli, R., & Hagmann, P. (2009). Predicting human resting-state functional connectivity from structural connectivity. PNAS, 106, 2035–2040. 10.1073/pnas.0811168106

Huang, G. B., Jain, V., & Learned-Miller, E. (2007). Unsupervised joint alignment of complex images. 2007 IEEE 11th International Conference on Computer Vision, 1–8. 10.1109/ICCV.2007.4408858

Jacobs, H. I. L., Muller-Ehrenberg, L., Priovoulos, N., & Roebroeck, A. (2018). Curvilinear locus coeruleus functional connectivity trajectories over the adult lifespan: a 7T MRI study. Neurobiology of Aging, 69, 167–176. 10.1016/j.neurobiolaging.2018.05.021

Jacobs, H. I. L., Becker, A., Sperling, R. A., Guzman-Velez, E., Baena, A., Uquillas, F. d’Oleire, … Quiroz, Y. T. (2019). Locus coeruleus intensity is associated with early amyloid and tau pathology in preclinical autosomal dominant Alzheimer’s disease. Alzheimer’s & Dementia, 15, P774–P775. 10.1016/j.jalz.2019.06.2832

James, T., Kula, B., Choi, S., Khan, S., Bekar, L., & Smith, N. (2021). Locus coeruleus in memory formation and Alzheimer’s disease. European Journal of Neuroscience, 54, 6948–6959. doi:10.1111/ejn.15045

Kellerman, T., Regenbogen, C., De Vos, M., Mößnang, C., Finkelmeyer, A. & Habel, U. (2015). Effective connectivity of the human cerebellum during visual attention. J Neuro, 33, 11453–11460. DOI: 10.1523/JNEUROSCI.0678-12.2012

Kerbler, G. M., Fripp, J., Rowe, C. C., Villemagne, V. L., Salvado, O., Rose, S., & Coulson, E. J. (2015). Basal forebrain atrophy correlates with amyloid β burden in Alzheimer’s disease. NeuroImage: Clinical, 7, 105–113. 10.1016/j.nicl.2014.11.015

Keren, N. I., Lozar, C. T., Harris, K. C., Morgan, P. S., & Eckert, M. A. (2009). In-vivo mapping of the human locus coeruleus. NeuroImage, 47, 1261–1267. doi: 10.1016/j.neuroimage.2009.06.012

Levitt, P. & Moore, R. Y. (1978). Noradrenaline neuron innervation of the cortex in the rat. Brain Research, 139, 219–231. 10.1016/0006-8993(78)90925-3

Liu, X., Ye, K., Weinshenker, D. (2015). Norepinephrine protects against amyloid-β toxicity via TrkB. J Alzheimers Dis, 44, 251–260. doi: 10.3233/JAD-141062

Madden, D. J. (2007). Aging and visual attention. Association for Psychological Science, 16, 70–74. doi: 10.1111/j.1467-8721.2007.00478.x

Madden, D. J., & Langley, L. K. (2003). Age-related changes in selective attention and perceptual load during visual search. Psychol Aging., 18, 54–67. doi: 10.1037/0882-7974.18.1.54

Markello, R. D., Spreng, N. R., Luh, W., Anderson, A. K., & De Rosa, E. (2018). Segregation of the human basal forebrain using resting state functional MRI. NeuroImage, 173, 287–297. 10.1016/j.neuroimage.2018.02.042

Mather, M., & Harley, C. W. (2016). The locus coeruleus: Essential for maintain cognitive function and the aging brain. Trends in Cognitive Science, 20(4), 214–227. 10.1016/j.tics.2016.01.001

Mesulam, M. M., Mufson, E. J., Levey, A. I., & Wainer, B. H. (1983). Cholinergic innervation of cortex by the basal forebrain: cytochemistry and cortical connections of the septal area, diagonal band, nucleus basalis (substantia innominata), and hypothalamus in the rhesus monkey. J Comp Neurol, 214, 170–197. doi: 10.1002/cne.902140206

Moyal, R., Turker, H. B., Luh, W., & Swallow, K. M. (2022). Auditory target detection enhances visual processing and hippocampal functional connectivity. Front. Psychol., 13, 10.3389/fpsyg.2022.891682

Munn, B. R., Müller, E. J., Wainstein, G., & Shine, J. M. (2021). The ascending arousal system shapes neural dynamics to mediate awareness of cognitive states. Nature Communications, 12, 6016. 10.1038/s41467-021-26268-x

O’Reilly, J. X., Croxson, P. L., Jbabdi, S., Sallet, J., Noonan, M. P., Mars, R. B., Browning, P. G. F., Wilson, C. R. E., Mitchell, A. S., Miller, K. L., Rushworth, M. F. S., & Baxter, M. G. (2013). Causal effect of disconnection of lesions on interhemispheric functional connectivity in rhesus monkeys. PNAS, 110, 13982–13987. 10.1073/pnas.1305062110

Peters, A., Setharas, C., & Luebke, J. I. (2008). Synapses are lost during aging in the primate prefrontal cortex. Neuroscience, 152, 970–981. doi: 10.1016/j.neuroscience.2007.07.014.

Riley, E., Turker, H., Wang, D., Swallow, K. M., Anderson, A. K., & De Rosa, E. (2023). Nonlinear changes in pupillary attentional orienting responses across the lifespan. GeroScience. 10.1007/s11357-023-00834-1

Sarter, M., Givens, B., & Bruno. J. P. (2001). The cognitive neuroscience of sustained attention: where top-down meets bottom-up. Brain Research Reviews, 35, 146–160. 10.1016/S0165-0173(01)00044-3

Shine, J. (2019). Neuromodulatory influences on integration and segregation in the brain. Trends in Cognitive Science, 23, 572–583. doi: 10.1016/j.tics.2019.04.002

Swallow, K. M., & Jiang, Y. V. (2010). The attentional boost effect: Transient increases in attention to one task enhance performance in a second task. Cognition, 115, 118–132. doi: 10.1016/j.cognition.2009.12.003

Swallow, K. M., & Jiang, Y. V. (2014). The attentional boost effect really is a boost: Evidence from a new baseline. Atten Percept Psychophys, 76, 1298–1307. doi: 10.3758/s13414-014-0677-4

Swallow, K. M., Broitman, A. W., Riley, E., Turker, H. B. (2022). Grounding the attentional boost effect in events and the efficient brain. Front Psychol., 13, 892416. doi: 10.3389/fpsyg.2022.892416

Taylor, N. L., D’Souza, A., Munn, B. R., Lv, J., Zaborsky, L., Muller, E. J., Wainstein, G., Calamante, F., & Shine, J. M. (2022). Structural connections between the noradrenergic and cholinergic system shape the dynamics of functional brain networks. NeuroImage, 260, 119455. 10.1016/j.neuroimage.2022.119455

Teipel, S., Heinsen, H., Amaro Jr., E., Grinberg, L. T., & Krause, B., Grothe, M. (2014). Cholinergic basal forebrain atrophy predicts amyloid burden in Alzheimer’s disease. Neurobiology of Aging, 35, 482–491. 10.1016/j.neurobiolaging.2013.09.029

Turker, H. B., Riley, E., Luh, W., Colcombe, S. J., & Swallow, K. M. (2021). Estimates of locus coeruleus function with functional magnetic resonance imaging are influenced by localization approaches and the use of multi-echo data. NeuroImage, 118047, doi: 10.1016/j.neuroimage.2021.118047.

Verissimo, J., Verhaeghen, P., Goldman, N., Weinstein, M., & Ullman, M. T. (2022). Evidence that ageing yields improvements as well as declines across attention and executive functions. Nature Human Behavior, 6, 97–110. 10.1038/s41562-021-01169-7

Wagatsuma, A., Okuyama, T., Sun, C. Smith, L., Abe, K., & Tonegawa, S. (2017). Locus coeruleus input to the hippocampal CA3 drives single-trial learning of a novel context. Proceedings of the National Academy of Sciences of the United States of America, 115, E310–E316. doi:10/1073/pnas.1714082115

Walling, S., Brown, R., Milway, J., Earle, A., & Harley, C. (2011). Selective tuning of hippocampal oscillations by phasic locus coeruleus activation in awake male rats. Hippocampus, 21, 1250–1262. doi:0.1002/hipo.20816.

Weinshenker, D. (2008). Functional Consequences of Locus Coeruleus Degeneration in Alzheimers Disease. Current Alzheimer Research, 5(3), 342–345. 10.2174/156720508784533286

Willenbockel, V., Sadr, J., Fiset, D., Horne, G. O., GOsselin, F., & Tanaka, J. W. (2010). Controlling low-level image properties: The SHINE toolbox. Behavior Research Methods, 42, 671–684. 10.3758/BRM.42.3.671

Yu, A. J., & Dayan, P. (2005). Uncertainty, neuromodulation, and attention. Neuron, 46, 681–692. Doi: 10.1016/j.neuron.2005.04.026

Zarow, C., Lyness, S. A., Mortimer, J. A., & Chui, H. C. (2003). Neuronal loss is greater in the locus coeruleus than nucleus basalis and substantia nigra in Alzheimer and Parkinson Diseases. Arch Neurol, 60, 337–341. doi: 10.1001/archneur.60.3.337

Zaborszky, L., Hoemke, L., Mohlberg, H., Schleicher, A., Amunts, K., & Zilles, K. (2008). Stereotaxic probabilistic maps of the magnocellular cell groups in human basal forebrain. NeuroImage, 42, 1127–1141. 10.1016/j.neuroimage.2008.05.055

